# Extinction and the temporal distribution of macroevolutionary bursts

**DOI:** 10.1101/725689

**Authors:** Stephen P. De Lisle, David Punzalan, Njal Rollinson, Locke Rowe

## Abstract

Phenotypic evolution through deep time is slower than expected from microevolutionary rates. This is the paradox of stasis. Previous models suggest stasis occurs because populations track adaptive peaks that typically move on million-year intervals, raising the equally perplexing question of why peaks shifts are so rare. Here, we consider the possibility that peaks can move more rapidly than populations can adapt, resulting in extinction. We model peak movement with explicit population dynamics, parameterized with published microevolutionary parameters. Allowing extinction greatly increases the parameter space of peak movements that yield the appearance of stasis observed in real data through deep time. Our work highlights population ecology as an important contributor to macroevolutionary dynamics, presenting an alternative perspective on the paradox of stasis where apparent constraint on phenotypic evolution in deep time reflects our restricted view of the subset of earth’s lineages that were fortunate enough to reside on relatively stable peaks.

## INTRODUCTION

Phenotypic change can be rapid at microevolutionary timescales (Hendry & Kinnison 1999; Kinnison & Hendry 2001; Gotanda *et al.* 2015), consistent with strong selection (Endler 1987; Conner 2001) and abundant genetic variance (Mousseau & Roff 1987). Yet, analyses of microevolutionary, fossil, and comparative data on body size reveal that divergence is unexpectedly modest on slightly longer timescales, with cumulative divergence becoming substantial only over macroevolutionary time (Estes & Arnold 2007; Uyeda *et al.* 2011). These new analyses indicate a protracted period of bounded evolution lasting for approximately one million years, prior to striking bursts of divergence in deeper time. Although similar patterns can be generated by a variety of macroevolutionary processes, statistical analysis indicates overwhelming support for a model of discrete, and rare, peak shifts to explain body size evolution (Uyeda *et al.* 2011; Arnold 2014). The pattern of limited phenotypic change over long timescales followed by macroevolutionary bursts suggests that some process must constrain phenotypic change within populations, as well as among closely related populations. Concomitantly, these results suggest that selection in the wild is net stabilizing, with shifts in the optimum phenotype occurring with exceptional rarity. Thus, rather than solve the paradox of macroevolutionary stasis, these new results instead highlight two fundamental challenges in reconciling micro and macroevolution: 1) if peak shifts are rare, as suggested by analysis of phenotypic data, do we see similar patterns of peak shifts in microevolutionary studies of selection in the wild, and 2) what processes may account for discrepancies in our inference of evolutionary dynamics across short and deep time? Identifying why phenotypic change is constrained over long timescales, and hence resolving the ‘paradox of stasis’, remains one of the most important problems to arise from the modern synthesis (Hansen & Houle 2004; Futuyma 2010; Arnold 2014; Pujol *et al.* 2018; Voje *et al.* 2018), embodying the fundamental challenge of identifying the microevolutionary processes that can explain patterns of divergence at greater timescales.

The only models that seem to explain body size evolution across timescales depend primarily on movement of an adaptive peak (or optimum), whereas genetic parameters play little explanatory role (Arnold 2014). In fact, the best-supported model is one of multiple peak displacements, where displacements of varying magnitude occur relatively infrequently (Uyeda *et al.* 2011). A problem that emerges when considering a model with peak movement is that, in real populations, displacements from the optimum phenotype will often be accompanied by reductions in mean absolute fitness and, thus, population size (Haldane 1937). In the extreme case, frequent or extreme displacements will result in an extinction event. Even under weak selection and modest but persistent peak movement, population size can fall below the replacement rate and the population will eventually go extinct (Lynch & Lande 1993; Bürger & Lynch 1995; Gomulkiewicz & Holt 1995). This suggests an important role for population ecology in macroevolutionary dynamics and highlights another unresolved question: what is the role of extinction in the observed distribution of phenotypic divergence in deep time? Although Uyeda et al. (2011) acknowledged that extinction could play a role generating the patterns observed in their data, they did not explicitly consider extinction in their likelihood-based approach to estimating model fits and parameters, or consider how extinction may influence estimates of peak shifts. A previous treatment (Estes & Arnold 2007) made similar restrictive assumptions regarding the magnitude of between-generation peak movement, on the grounds that displacements of excessively large magnitudes would result in the extinction of lineages. Yet the importance of extinction for patterns of phenotypic diversity are broadly recognized (Jablonski 2008), and moreover, the observation that the vast majority of biodiversity is now extinct (van Valen 1973) suggests extinction has played a critical role in shaping the patterns of diversity we see both today and in the past.

In the present study, we attempt to reconcile patterns of morphological stasis with observed variation in selection in the wild, through the application of macroevolutionary models that explicitly incorporate selection and population dynamics. Our goal was not to reanalyze the previous studies or critique any specific model of evolution, but rather highlight the effects of a general feature of biological populations, extinction, that has rarely been considered in models of phenotypic macroevolution. We focus on a subset of models of peak movement previously identified as the prime candidates for the observed data (Arnold 2014). We simulated evolution using empirically-derived estimates of peak displacement, while simultaneously considering the effect of these displacements on the likelihood of extinction by tracking population dynamics explicitly. Although non-random extinction is known to influence our ability to interpret macroevolutionary models (Maddison 2006), the potential effects of extinction on our ability to estimate the dynamics of the adaptive landscape are currently unclear. We show that 1) observed direct estimates of fluctuations in the optimum phenotype in wild populations can be high, indicating that changes in the optimum are far more commonplace and severe than inferred from previous analyses of phenotypic change, and 2) incorporating extinction explicitly into macroevolutionary models of peak movement indicate that lineage loss can contribute substantially to apparent patterns of morphological stasis. Our work suggests that inference on the movement of adaptive peaks using observed phenotypic data alone fail to capture the fact that lineage loss may erase the history of rapid or severe peak shifts. Moreover, our work demonstrates that explicit integration of population ecology may shed light on patterns of phenotypic evolution in deep time.

## METHODS

### Basic approach to simulating phenotypic evolution and population size

We simulated the evolution of a quantitative trait in replicate populations, where populations were subject to several different scenarios determined by a range of evolutionary genetic parameters. We assumed phenotypic selection acting on a single, continuously distributed trait, *z*, with a population mean phenotype, 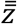, experiencing selection approximated by a Gaussian fitness function with a width, ω, with the position of a single optimum located at θ, and described by:

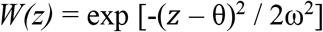

(see (Lande 1979; Estes & Arnold 2007)). We simulated evolution of 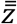 for up to 100,000 generations while allowing θ to vary in position on a generation-by-generation basis (i.e. assuming discrete time with non-overlapping generations). The behavior of θ was governed by processes simulating either Brownian motion of the optimum or peak displacement with (potentially) multiple bursts (see Uyeda *et al.* 2011 and below). In addition to tracking phenotypic evolution, at every generation, *t*, we allowed population size (*N*) to change according to the average fitness of the population, 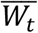, which depends on the phenotypic distribution relative to the optimum following

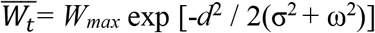

where *W_max_* corresponds to maximum absolute fitness, *σ*^2^ equals the phenotypic variance, and *d*^2^ is equivalent to 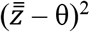^2^ (Gomulkiewicz & Holt 1995, Eqn. 6; Estes & Arnold 2007, Eqn. 2).

To model the more realistic scenario of density-dependent population growth associated with phenotypic evolution, we assumed (logistic) population growth, *r*, is a function of the maximum growth rate ln(*W*_max_) and the distance to the new optimum, described by:

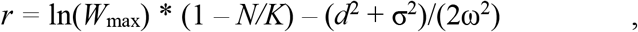

after Lynch and Lande (1993, Eqn 2), and incorporating load on population growth introduced by phenotypic variance (*σ*^2^) (see Kirkpatrick and Barton (1997, Eqn. 7)). We assumed that, under a given selection scenario (i.e. for all replicates for a set of parameters, for the duration of the simulated interval) the shape of the fitness function remained constant. In the models considered, the response to selection depended on the available genetic variance (expressed as heritability), *h^2^*.

We focused on two classes of models of peak movement: one that invokes movement of the optimum at relatively constant rate: the Brownian motion of the optimum (Felsenstein 1988) and another that envisions more sporadic movement: the peak displacement or ‘burst’ model (sensu Estes & Arnold 2007; Uyeda *et al.* 2011). To estimate the impact of extinction, we contrasted each model of peak movement when extinction was allowed and when extinction was prevented. In our simulations, extinction was prevented by simply re-seeding the population with some minimum *N* if the population size fell to zero, or below some critical threshold.

Although past workers have made use of explicit likelihood functions for alternative models of peak movement that allow comparisons with observed data (Estes & Arnold 2007; Uyeda *et al.* 2011), we aim to explore the qualitative effects of peak movement on population dynamics to examine different scenarios that succeed or fail to generate the characteristic pattern of relative stasis, followed by bursts of change (i.e. qualitatively resembling the ‘blunderbuss’ pattern observed by Uyeda *et al.* 2011). In the models of Brownian motion of the optimum, at each time step (*t* + 1), the position of θ was determined by its position at *t*, plus a deviation with expected mean = 0 and variance = σ_θ_^2^. Brownian motion models with (BME) and without extinction (BM) differed only in that the latter allowed the populations to be rescued whenever the population fell below *N* = 50.

To model a scenario where the optimum experiences displacement less frequently than at every generation, the value of θ at a given time step was determined by its previous position but potentially also by a displacement that occurred with some probability. As in Uyeda et al. (2011), the probability of a displacement event occurring was modeled as a Poisson process, determined by the parameter, *λ*, or the average expected number of displacement events per generation. In instances where a displacement event occurred, its magnitude was drawn from a normal distribution with mean = 0 and variance = σ_θ_^2^. As above, displaced optimum models with (DOE) and without extinction (DO) were distinguished by allowing the latter populations to be rescued whenever *N* < 50. Note that both the probability of a peak displacement λ and variance of peak displacements σ_θ_^2^ together determine the total rate of peak movement in this model. The distribution of peak movements after a given amount of time is N(0, mσ_θ_^2^), where m is the expected number of peak displacements (Uyeda *et al.* 2011). As such, peak shifts that are large but extremely rare result in a slow average rate of peak movement through time, while frequent peak shifts of moderate magnitude would result in a higher average rate of peak movement.

### Model parameters

Whenever possible and appropriate, we used parameter values that are similar or the same as in past work (Estes & Arnold 2007; Uyeda *et al.* 2011). One major difference, however, is that we used a dataset of temporally replicated estimates of phenotypic selection compiled by Siepielski et al. (2011) to calculate the empirical distribution of displacement distances. For all studies in the database that recorded both standardized linear, β, and negative nonlinear (corresponding to approximately stabilizing) selection gradients, γ, we calculated the distance to the optimum within each selective episode as:

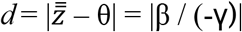

(Phillips & Arnold 1989, Eqn 11; Estes & Arnold 2007, Eqn 7). By convention, gradients were normalized to a mean of 0 and σ = 1 (Lande & Arnold 1983). The standardized selection gradients allow estimation of the relative distance between 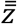 and θ within a given *i*^th^ temporal replicate or ‘episode’, but provide no direct information on the actual values/coordinates of θ. Nonetheless, under the assumption that at the *i*^th^ episode, 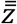 is closer to its optimum than it was at episode *i* - 1, the difference in absolute distances, Δ*d*, between a given pair of successive replicates provides a proxy for the total displacement of the optimum between episodes. The variance of the distribution of Δ*d*, σ_θ_^2^, was used to parameterize peak movement in the simulations, assuming a normal distribution of possible displacements as in the burst models of Uyeda et al. (2011). We relaxed this assumption of normality by also running the same models but with peak displacements drawn from the empirical estimates of Δ*d*. Implicit in our use of the empirical data to approximate displacements of θ is that variation in selection reflects movement of the optimum, and not temporal variability in the phenotypic distribution. We recognize that the Siepielski et al. (Siepielski *et al.* 2009) dataset may reflect both sources of variation and results in unbiased, albeit imperfect, estimates of peak displacements. Following Estes and Arnold’s (Estes & Arnold 2007) treatment of similar data, we excluded several observations corresponding to questionably large values of θ (i.e. > |35| standard deviations) that most likely reflect gradients associated with large estimation errors.

Each simulation began (i.e. at *t* = 0) with a population mean phenotype, 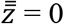, and phenotypic variance, σ_*p*_^2^ = 1. We arbitrarily set *W_max_* = 1.5, which corresponds to rapid exponential population growth (Malthusian *r* = 0.41) for populations residing at the optimum. Variation in *W_max_* had little effect on conclusions under density dependent population growth, as this growth rate is rarely experienced (e.g., a small population residing at its optimum phenotype). For each simulated population, carrying capacity (*K*) was randomly drawn from a range of 10561 < *K* < 12259, based on estimates for vertebrates (see Table 2 in Reed *et al.* 2003), and each simulation began with a population at carrying capacity (i.e. *N* = *K*). For each population, heritability (*h^2^*) was drawn from another empirical dataset on vertebrate body size (Hansen *et al.* 2011), and we assumed a constant *h^2^* throughout a given simulation. For all simulations, we assumed an adaptive landscape with a width of ω^2^ = 3, comparable to median values from previous synthetic studies (Estes & Arnold 2007). Although we did not exhaustively explore the entire parameter space, the qualitative results of the simulations (i.e. the shape of the plots) were generally robust to changes in parameters, though notable exceptions and their interpretation are addressed in the Results and Discussion. For each model, we simulated 500 replicate lineages for up to 100,000 generations. This interval is several orders of magnitude shorter than the timescale considered in Uyeda et al. (2011), but was necessitated by computational limitations, including the graphical requirements of plotting phenotypic trajectories. Similarly, our scale of divergence (in units of phenotypic standard deviation) differs from the one used by Uyeda et al. (2011), who reported divergence in units of log-differences in body size. However, our main objective was to reach general conclusions regarding factors that influence the *relative* temporal distribution of evolutionary change, and this is not considered to be strongly dependent on the interval or scale (Uyeda *et al.* 2011). For example, the waiting time for the ‘burst’ divergence in our displaced optimum models were allowed to occur sooner by setting larger values of λ, without any complications in interpretation. The expected distribution of phenotypic divergence under a multiple burst model is N(0, *m*σ_θ_^2^), and the number of bursts *m* in a given timescale *t* is given by m = ∑_*t*_ *λ*, and as such, performing simulations over a shorter timescale *t* under greater *λ* allows for the analysis of similar total evolutionary divergence (number of bursts) as the empirical data. Finally, we compared the distributions (variances) of phenotypic divergence at the end of each simulation run using Levene’s test for equality of variances, and we also plot the accrual of phenotypic variance through time. Simulations were implemented using the R base package (R Core Team 2017), and examples of code for our 4 main classes of models (BM, BME, DO, DOE) are available in the Supplementary Material, S1. The subsets of data analyzed to generate estimates of Δ*d* and from which *h^2^* was sampled are available on Dryad.

## RESULTS

Our analysis of the empirical dataset of temporally-replicated selection gradients resulted in 369 estimates of θ from 138 studies. From these we obtained 231 estimates (from 87 studies) of temporal displacements of the optimum, Δ*d*. The distribution of displacements (around zero) of the optimum was roughly symmetric (Figure 1) around zero, with a Laplace distribution with a standard deviation of 12.98. Although the median magnitude (|Δ*d*|) was 1.21σ (mean = 3.92σ, standard error = 0.82) is comparable to the estimates for θ in Estes and Arnold (2007), our empirical estimate of the standard deviation of Δ*d* is far higher than estimates of the variation in Δ*d* estimated in past modelling efforts. Specifically, ML estimates of the magnitude of peak displacement from ref 8 correspond to a standard deviation of Δ*d* of approximately 3 phenotypic standard deviations (Arnold 2014), an order of magnitude less than the empirically-observed value.

**Figure 1.**
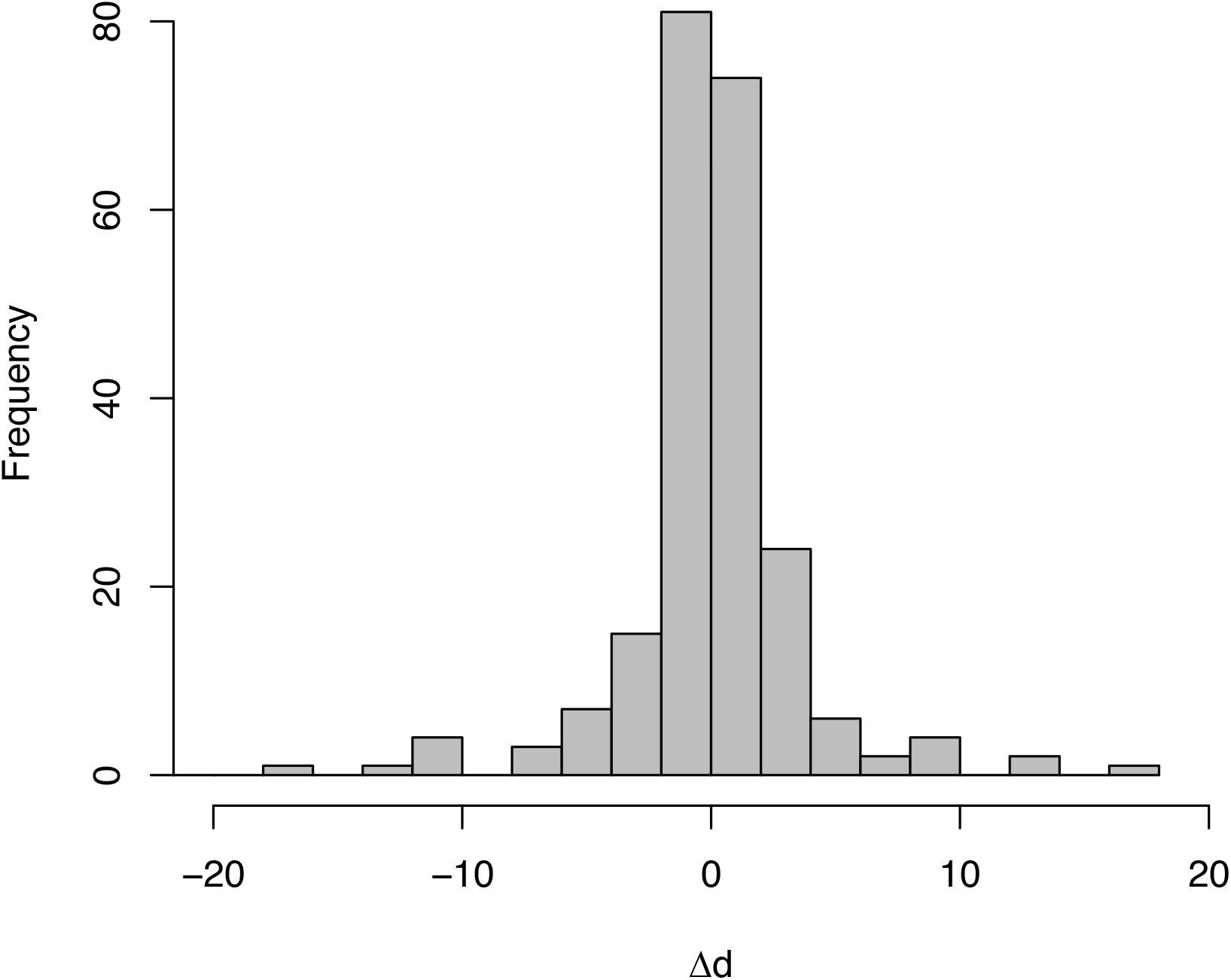
Histogram of estimated displacements, Δ*d*, of the optimum based on the dataset of Siepielski et al. (Siepielski *et al.* 2009). To aid in visualization, estimates of |Δ*d*| > 20 are excluded.

Our BM and BME simulations underline two ways in which Brownian motion models fail to capture empirical patterns at both the micro and macro scale. At a low value of σ_θ_ = 0.1, the pattern of replicated divergence is confined to a narrow band that persists long enough to appear as a period of initial stasis. However, a moderate increase of σ_θ_ to 0.5 results in phenotypic divergence that is too rapid, and inconsistent periods of protracted stasis. These patterns appear insensitive to whether or not extinction is permitted when examining plots of divergence through time (Fig. 2a, b, left panels). However, allowing extinction decreases the extent of phenotypic divergence in deep time, and this effect increases with increasing σ_θ_^2^ (Levene’s test for σ_θ_ = 0.5: *F*_1,998_ = 447, *P* < 2.2*10^−16^, Fig. 2B right panel; Levene’s test for σ_θ_ = 0.1: F1,998 = 2.04, *P* = 0.15, Fig. 2A right panel; See also Figure 3). Arming the BM models with our empirical estimate of the variance in Δ*d* generates two opposing and equally unrealistic patterns of divergence. When extinction is not allowed, the frequent and persistent movements of the peak result in rapid divergence that almost immediately spans the entire phenotype space; when extinction is permitted, such peak movement results in only moderate divergence before all lineages rapidly go extinct (Figure 2c). Thus, our results not only suggest BM models fail in that they result in too much divergence under realistic parameter values, as noted in the past (Conner 2001; Arnold 2014), but also that these models fail even with the inclusion of extinction because peak movement can be rapid enough that nearly all lineages fail to survive.

**Figure 2.**
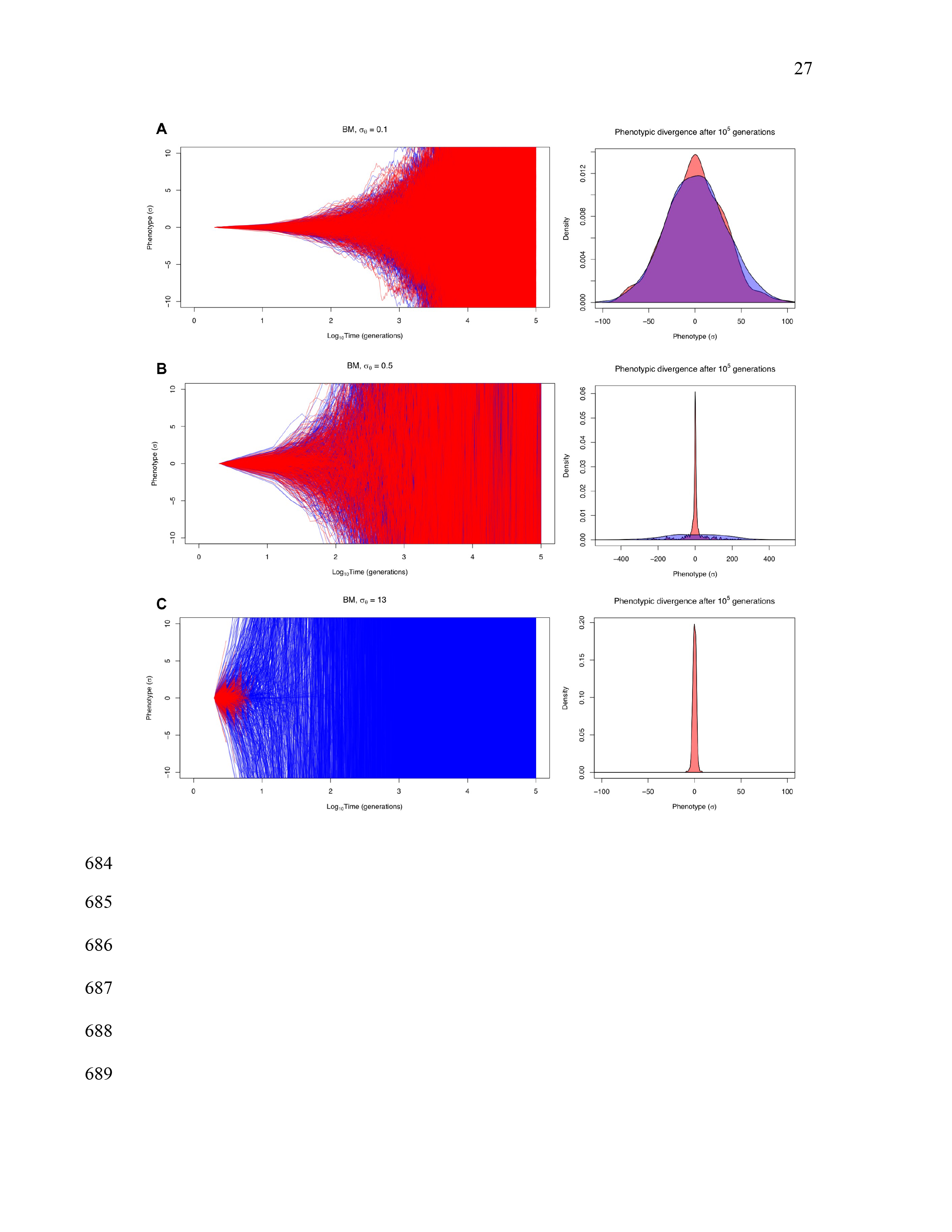
Brownian motion models (BM, blue; BME, red) when altering the rate of peak movement, σ_θ_. BME models allow the possibility of extinction for maladapted populations, while populations in BM simulations are ‘rescued’ from potential extinction (see text). A and B represent two low-moderate values of σ_θ_^2^, while C assumes σ_θ_^2^ derived from empirical estimates from wild populations. Right panels indicate the phenotypic distributions at the end of the simulations; either at extinction or 10^5^ generations. Note scale differences in x axes of right panels.

**Figure 3.**
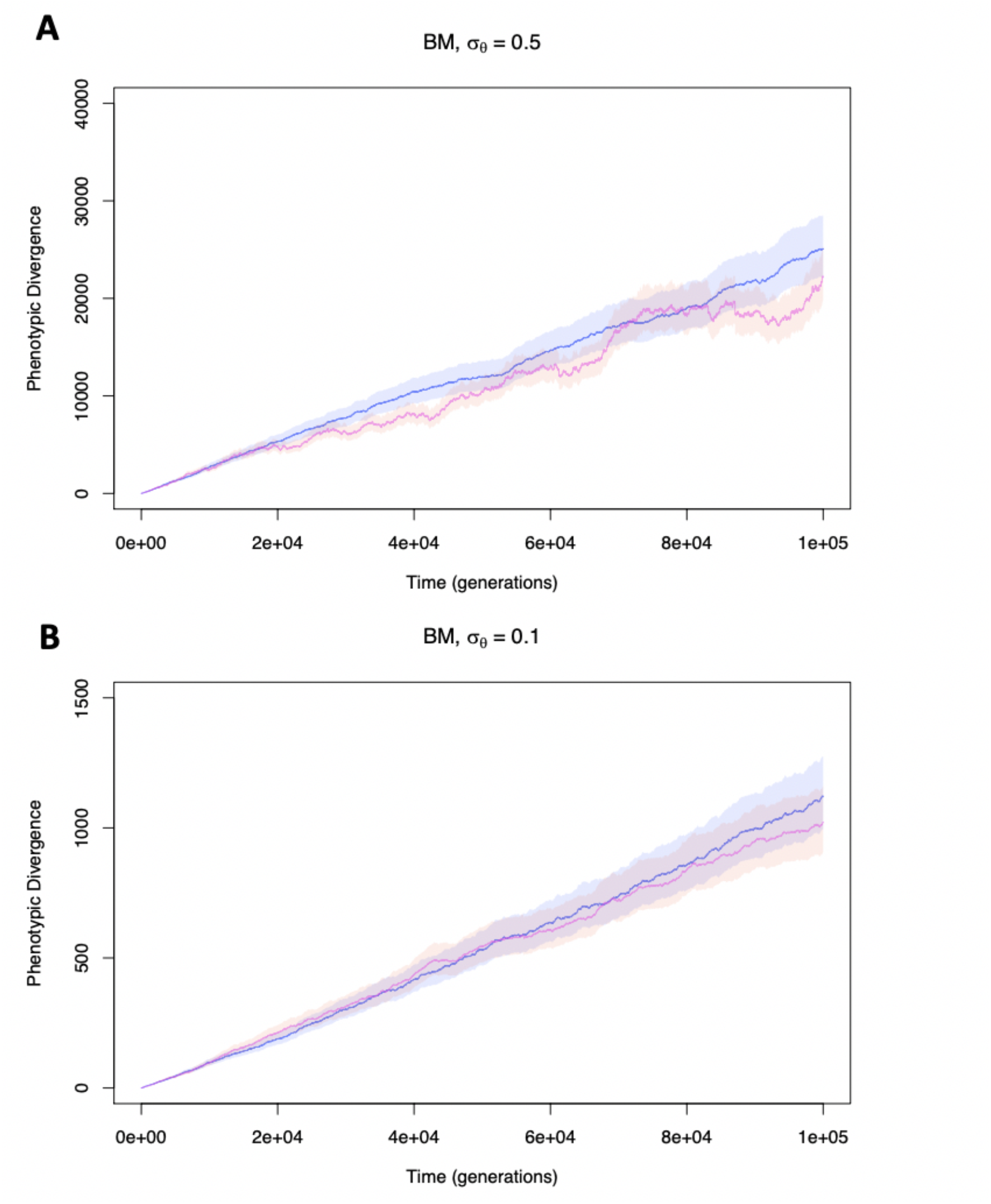
Variance accrual through time in Brownian motion models (BM, blue; BME, red) when altering the rate of peak movement, σ_θ_^2^. BME models allow the possibility of extinction for maladapted populations, while populations in BM simulations are ‘rescued’ from potential extinction (see text). A and B represent two low-moderate values of σ_θ_^2^.

Next, we modeled discrete peak displacements that occur with varying frequency, from extremely infrequent (i.e. *λ* = 10^−6^, which approaches the rate of approximately 1 in 10^7^ estimated by Uyeda et al. (2011)), to two orders of magnitude higher (*λ* = 10^−4^) but with the same potential magnitude. This case yields a pattern resembling that empirically observed by Uyeda et al. (2011), but the narrow band of initial divergence becomes considerably more distinct when extinction is permitted (Figure 4, left panels). When peak shifts are extremely rare over the timescale of simulations, little phenotypic divergence is observed (Fig. 4A), although even in this case extinction has substantial consequences on the amount of divergence observed in deep time (Levene’s test: *F*_1,998_ = 7.85, *P* = 0.005, Fig. 4A right panel). Increasing the frequency of peak shifts by an order of magnitude has severe demographic consequences, in turn restricting early divergence (Fig. 3b, left panel), and illustrating the potential for extinction to generate a pattern of apparent evolutionary constraint (Levene’s test: *F*_1,998_ = 234, *P* < 2.2*10^−16^, Fig. 4B, right panel). When we assume that peak displacements still occur sporadically but with even higher frequency (*λ* = 10^−4^), some key insights are revealed. Extinction becomes critical to shaping the temporal pattern of phenotypic evolution, as divergence becomes otherwise unconstrained over longer timescales (Levene’s test: *F*_1,998_ = 737, *P* < 2.2*10^−16^, Fig. 4C, right panel). Shorter waiting times between displacements lead predictably to a shortening of the period of relative stasis, and a shorter average longevity of lineages (Figure 4c, right panel). More frequent displacement events also result in a coarse transition between the period of stasis to the eventual ‘bursts’ of evolution, implying that longer periods of favorable conditions, and therefore more stable population sizes, are more conducive to the formation of patterns to those that have been empirically observed^8^. The substantial effect of including extinction can be readily observed in the accrual of phenotypic variance through time, which is drastically reduced when extinction is permitted (Fig. 5A-C), even when peak shifts are exceptionally rare (*λ* = 10^−7^) and the distribution of Δ*d* is reduced to 3 phenotypic standard deviations (Fig. S3).

**Figure 4.**
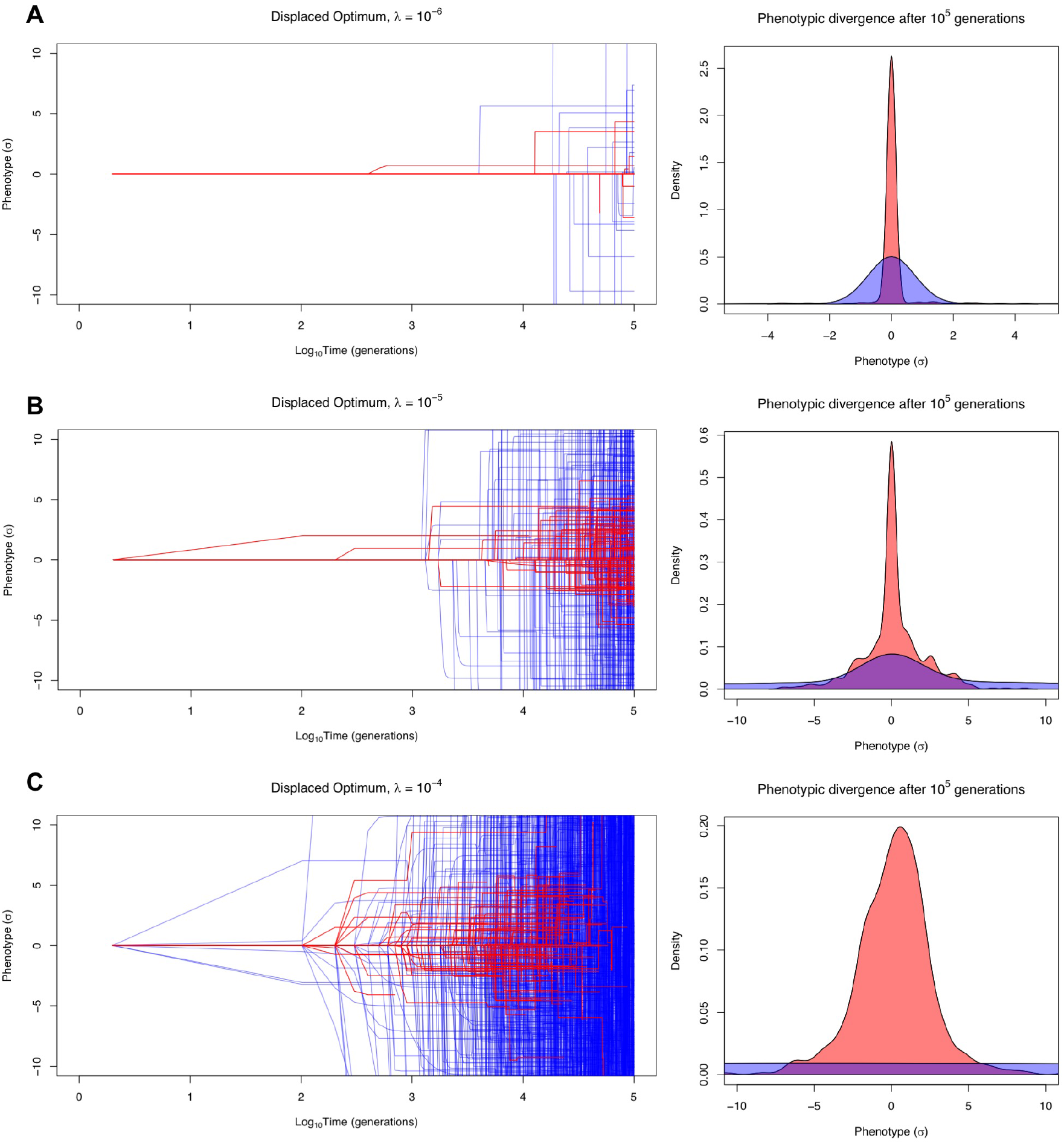
Displaced optimum models (DO, blue; and DOE, red), assuming σ_θ_ = 13 (the empirically observed value) at different assumed probabilities of a peak shift, *λ*. DOE models allow the possibility of extinction for maladapted populations, while populations in DO simulations are ‘rescued’ from potential extinction (see text). Right panels indicate the phenotypic distributions at the end of the simulations; either at extinction or 10^5^ generations. Note scale differences in x axes of right panels.

**Figure 5.**
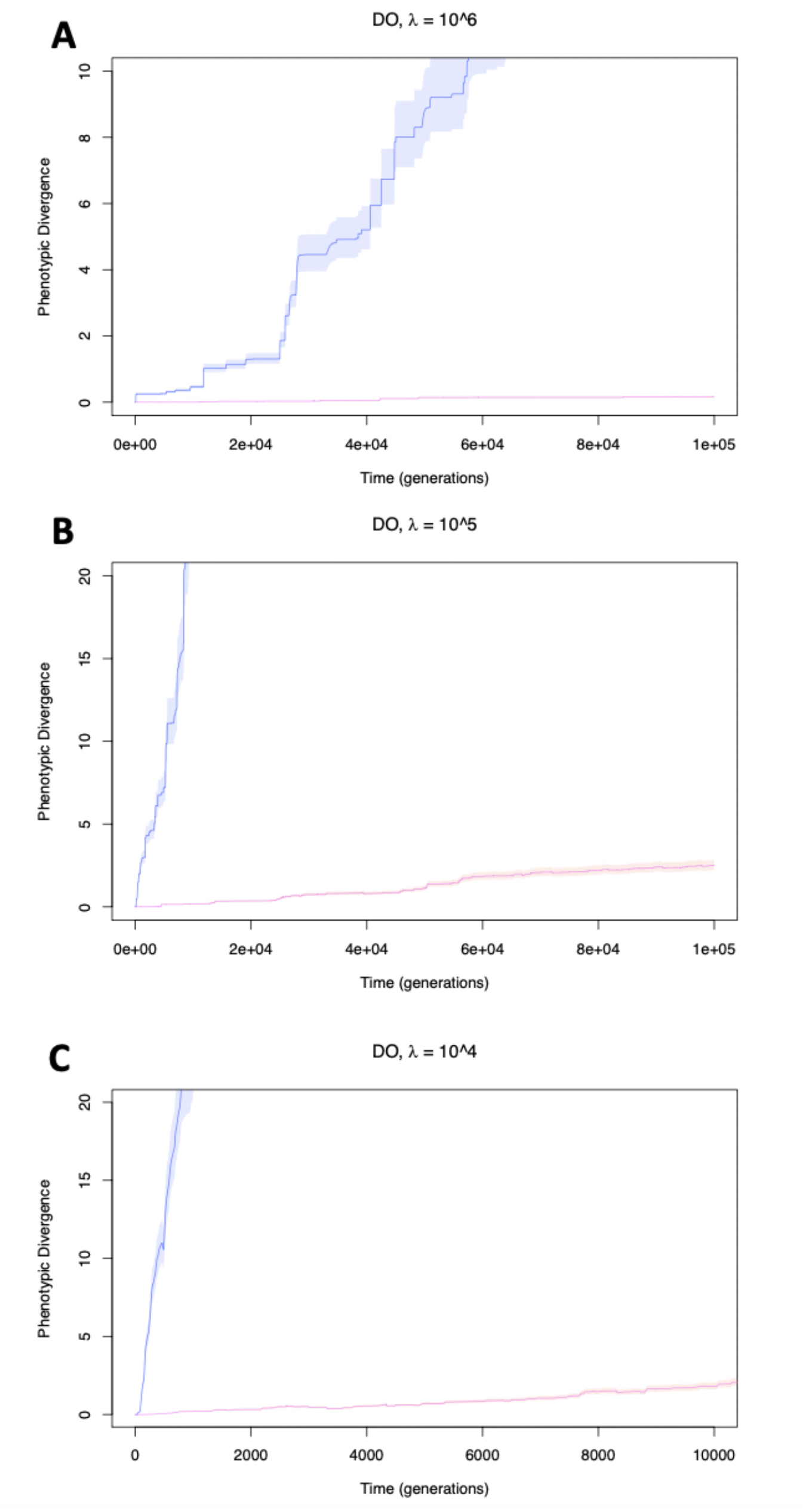
Variance accrual through time in displaced optimum models (DO, blue; and DOE, red), assuming σ_θ_ = 13 (the empirically observed value) at different assumed probabilities of a peak shift, *λ*. DOE models allow the possibility of extinction for maladapted populations, while populations in DO simulations are ‘rescued’ from potential extinction (see text).

Our conclusions were essentially unchanged when re-running the model with displacements randomly drawn from the (exponential) empirical distribution of Δ*d* instead of a normal distribution (Supplementary Material, Figure S2).

Although the empirical estimates used to parameterize σ^2^_θ_ may be biased upwards by sampling error, equivalent conclusions are obtained assuming a value σ^2^_θ_ that is an order of magnitude lower than the observed value (Levene’s test, *F*_1,998_ = 78, *P* < 2.2*10^−16^), and the accrual of variance through time is significantly reduced by extinction at long timescales even when σ_θ_ is reduced to 3 sds (Fig. S3). Only when decreasing σ^2^_θ_ by two orders of magnitude, to the order of a single phenotypic standard deviation, does the effect of extinction on macroevolutionary divergence disappear (Levene’s test, *F*_1,998_ = .98, *P* = .32). Yet such a small value of σ^2^_θ_ is inconsistent with the empirical observation that the divergence is confined to an interval that is several within-population phenotypic standard deviations wide (Uyeda *et al.* 2011; Arnold 2014). Thus, our conclusions regarding the importance of extinction in shaping macroevolutionary patterns appear robust to any potential biases in our empirical estimate of σ^2^_θ_.

Altering the form of population growth further reveals the importance of population size for withstanding temporal displacements, and ultimately, the accrual of phenotypic change. Substituting density-dependent growth with density-independent geometric growth (e.g., following Gomulkiewicz & Holt 1995, Eqn 6), using the same generous value of *W_max_* = 1.5 corresponds to unbounded, exponential population growth when populations reside at the optimum. Under such permissive growth conditions, the pattern of divergence remains similar so long as extinction is allowed, but much more divergence accumulates as a consequence of lower extinction rates and longer-lived lineages (Supplementary Material, Figure S3). This is attributable to the fact that prolonged periods of stability allow a population to grow to an extremely large size, which buffers the population from subsequent peak movement. Collectively, these results implicate limits to population persistence as a key factor in generating observed patterns of phenotype evolution, and more generally the results underscore the importance of demographic considerations for generating macroevolutionary patterns.

## DISCUSSION

Our analyses illustrate an important and previously unexplored role for extinction and population dynamics in generating patterns of apparent constraint in macroevolution. We show that in the presence of rapidly moving optima, populations may face extinction while attempting to track their adaptive peak, preventing these rapid peak shifts from being recorded as phenotypic change. Our work suggests that the observation of morphological stasis over macroevolutionary time may not reflect a lack of movement in phenotypic optima, but instead reflect our censored view of phenotypic evolution, which is restricted to those lineages that have not gone extinct. Observed patterns of phenotypic divergence in a variety of taxa (Estes & Arnold 2007; Uyeda *et al.* 2011; Arnold 2014) can be partly accounted for by an accrual of changes in many long-lived lineages fortunate to have experienced only small peak displacements, combined with a subset of lineages that, due to abrupt displacements, went extinct before much phenotypic divergence had accumulated.

Using a dataset of temporally-replicated estimates of phenotypic selection from the wild, we show that the distribution of peak shifts in extant natural populations is inconsistent with the rarity of such events as inferred from analysis of observed macroevolutionary phenotypic change, providing quantitative support to the idea that these past results are indeed paradoxical (as suggested by Uyeda *et al.* 2011). Concomitantly, such frequent and extreme shifts lead to predicted macroevolutionary rates that are far more extreme than those observed in empirical data. However, incorporating extinction into these macroevolutionary simulations results in temporal patterns of phenotypic divergence similar to empirical observations (e.g., the blunderbuss), and in all cases resulted in a decrease in the variance in phenotypic change observed in deep time (Figs. 2, 3, right panels). This suggests non-random extinction may play a key role in resolving the stasis paradox; although we frequently observe substantial peak shifts in extant populations (Feya *et al.* 2015), many of these populations would be expected to face extinction over longer timescales in the face of such maladaptation, leaving a pattern of apparent morphological stasis over macroevolutionary time despite strong selection observed in extant populations. The relative insensitivity of our results to assumptions of heritability further suggest that extinction can play an important role in patterns of long-term phenotypic evolution, even when selected traits are evolvable.

Our results are consistent with the observation that extinction is an important contributor to the distribution of Earth’s diversity, as most of life that has existed has gone extinct, and suggests that considering extinction events may be critical to understanding shifts in phenotypic optima through the history of life. The extent to which ignoring extinction will impact estimates of peak movements is illustrated in the effects extinction has on the observed variance in phenotypic change (Fig. 2, 4, left panels, Fig. 5). Because both Brownian Motion and Displaced Optimum models rely on the observed distribution of phenotypic divergence in deep time to obtain estimates of peak movement, where the distribution is treated as N(0, tσ_θ_^2^) (BM) or N(0, mσ_θ_^2^) (DO), any effect of extinction on the realized distribution of phenotypic change, if left unaccounted for, will be borne out in parameter estimates for peak movement that are lower than reality. Nonetheless, our results suggest the importance of extinction on the inference of peak shifts will depend upon the value of σ_θ_^2^. We have used larger values than previous workers (Estes & Arnold 2007; Uyeda *et al.* 2011). Yet as suggested by empirical data presented here as well as the frequency of directional selection in the wild (Kingsolver *et al.* 2001), large and frequent shifts in trait optima appear to be a reality of natural populations. Our study suggests that such shifts in the optimum phenotype observed in extant populations are compatible with the observation of macroevolutionary stasis, if the probability of extinction increases with the magnitude and frequency of peak shifts. The observation that mass extinctions correlate with geologic periods characterized by environmental upheaval (which would be expected to reflect rapid changes in optimal phenotypes), including current human-caused extinctions of the Anthropocene, further supports our conclusion that extinction may play a role in generating apparent morphological stasis.

Extinction will contribute to the appearance of stasis whenever it scales with the frequency or magnitude of peak shifts. Compared to the demographic rescue model in which extinction does not occur, models allowing extinction consistently result in a delay in phenotypic divergence, conferring a pattern that is visually distinct from the immediate divergence observed in the BM models. This pattern appears to hold under a wide range of values of stabilizing selection (1.5 > ω^2^ > 20), though very strong curvature severely restricts evolution to a narrow range for the entire ‘lifetime’ of the lineage (results not shown). Similarly, varying the range of heritability did not qualitatively change the importance of extinction. For example, the effect of decreasing heritability from *h^2^* = 0.4 to *h^2^* = 0.1 was, as expected, primarily to reduce the extent of divergence at all timescales. We also incorporated a “white noise” parameter in a subset of our simulations (not shown), in the same manner described by Estes and Arnold (Estes & Arnold 2007). This too had no impact on our conclusions. Thus, a role for extinction in generating patterns of apparent stasis do not appear to be qualitatively dependent on values of heritability, the curvature of the fitness surface, or white noise. The relatively small role of these parameters for explaining broad patterns is consistent with the findings of Estes and Arnold (Conner 2001).

Much of the phenotypic diversity that has ever emerged was probably quickly lost, and extinction has long been recognized as a potentially important force in phenotypic macroevolution; for example, explaining apparent disparities in speciation rates estimated for different timescales in (the ‘Ephemeral Speciation Model’; Futuyma 1987; Rosenblum *et al.* 2012; Auilée *et al.* 2018). Our study complements this view, but with an important distinction: our models do not invoke speciation and only permit anagenetic change. While the Ephemeral Speciation model emphasizes divergence that was aborted, our models emphasize the importance of extinction for the divergence that cannot accrue in the first place, representing simply one example of the general challenge that extinction imposes for comparative biology. As our ability to infer microevolutionary fitness surface is limited to the distribution of phenotypes we observe, our understanding of the shape and dynamics of macroevolutionary adaptive landscapes is limited by a reliance upon those lineages that have not gone extinct. Thus, extinction poses a challenge for all comparative methods that exploit phenotypic variation to infer nuances of the adaptive landscape, or even estimate rates of phenotypic evolution. Although this issue of extinction has been made especially clear for evolutionary rates of discrete traits (Maddison 2006), it seems especially germane when the very goal of the evolutionary model is to make a statement on (mal)adaptation (Hansen 1997; Bartoszek *et al.* 2012) or the dynamics of macroevolutionary adaptive peaks (Uyeda & Harmon 2014). The only potential resolution to the challenge extinction poses to comparative biology is to incorporate, whenever possible (e.g., Maddison *et al.* 2007), the dynamics of lineage accumulation explicitly into macroevolutionary models of trait evolution. Our simulation models represent a first step in this effort to estimate peak movement. Future work further integrating selection, population and macroevolutionary dynamics may present an important way forwards in resolving the paradox of macroevolutionary stasis.

## ACKNOWLEDGEMENTS

We are grateful to Erik Svensson and John Waller for discussion, and Joseph Uyeda for thoughtful criticisms and comments that improved the manuscript. Funding for this work was provided by grants from NSERC and the Canada Research Chairs Program to Locke Rowe. DP and NR also benefitted from the National Institute for Mathematical and Biological Synthesis, Evolutionary Quantitative Genetics Tutorial (2014).

## SUPPLEMENTARY MATERIAL

**S2.**
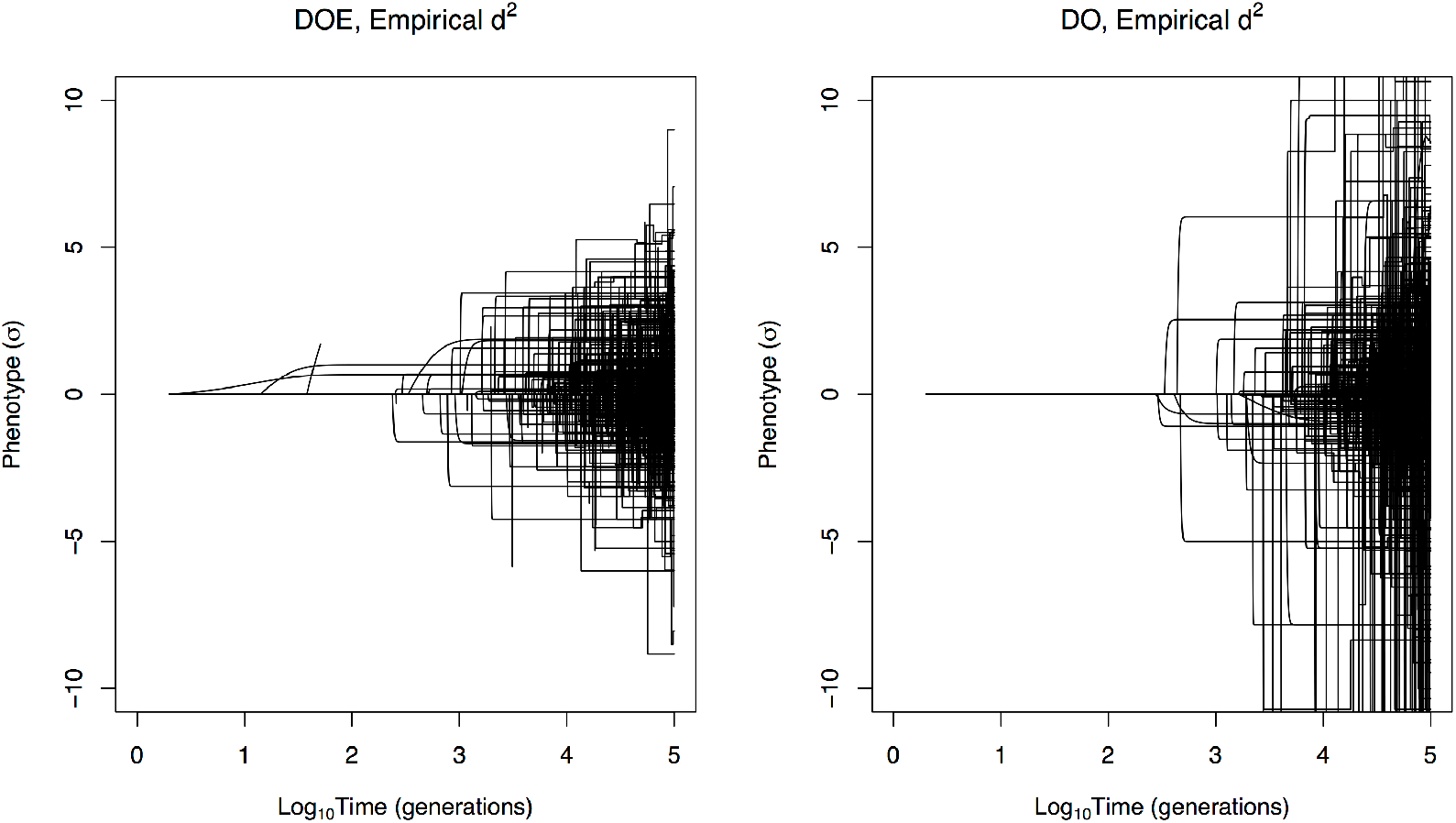
Phenotypic evolution in Displaced Optimum models with (DO, right panel) and without extinction (DOE, left panel) assuming *λ* = 10^−5^, with σ_θ_^2^ determined by drawing from the empirical distribution of Δ*d.* All other parameters were the same as those in Figure 3A.

**S3.**
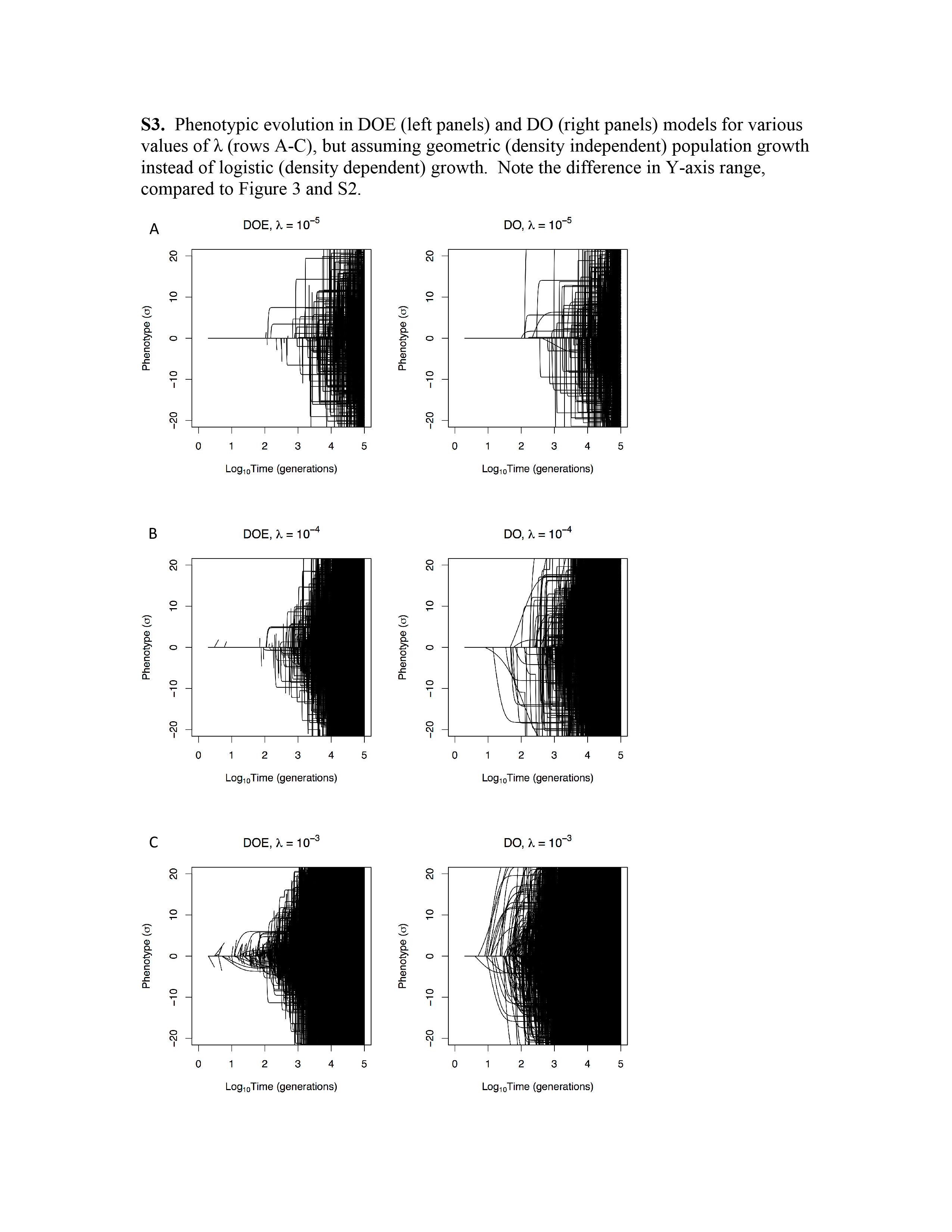
Phenotypic evolution in DOE (left panels) and DO (right panels) models for various values of *λ* (rows A-C), but assuming geometric (density independent) population growth instead of logistic (density dependent) growth. Note the difference in Y-axis range, compared to Figure 3 and S2.

**Figure S3.**
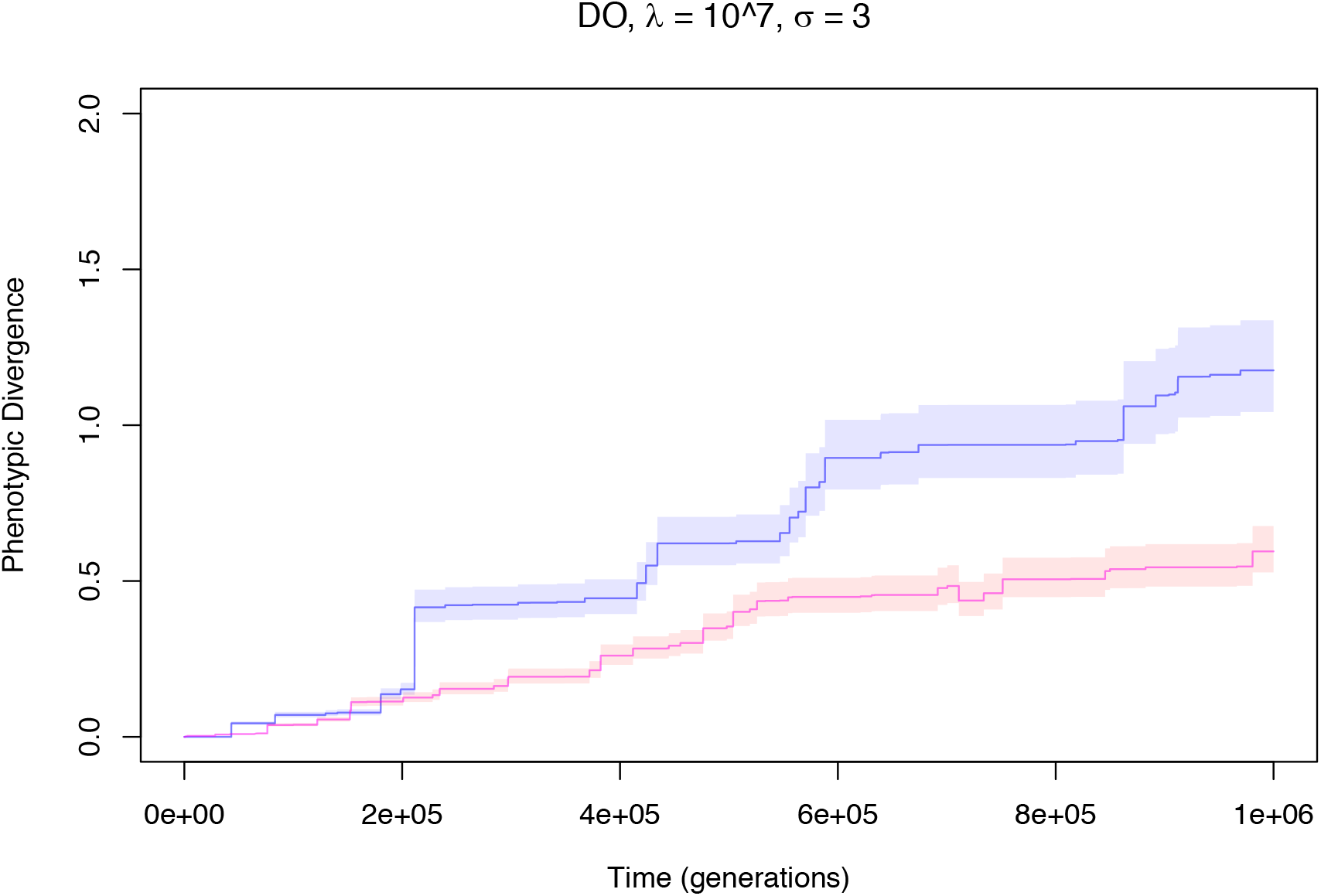
Variance accrual through time in displaced optimum models (DO, blue; and DOE, red), assuming σ_θ_ = 3 and extremely low probability of a peak shift, *λ*. DOE models allow the possibility of extinction for maladapted populations, while populations in DO simulations are ‘rescued’ from potential extinction (see text).

